# Steady state flux balance analysis game

**DOI:** 10.1101/2021.11.02.466952

**Authors:** Garud Iyengar, Mitch Perry

## Abstract

Flux balance analysis (FBA) for microbial communities often assumes a global objective function that all species cooperatively maximize in addition to maximizing their own growth. Combining community FBA with dynamic FBA to understand the time course and steady states of communities typically entails discretizing time and solving a community FBA model at each time point, a time-intensive process. We propose a dynamic community FBA model where species compete for metabolites to grow off of without needing to cooperate to maximize a community-level objective. An efficient method for computing steady state community compositions is provided, as well as methods for determining the stability of a steady state community to perturbations in biomass and invasion by species outside the community. The model and methods are applied to a model of four *E. coli* mutants with elements of competition (for shared metabolites) and cooperation (via mutants being auxotrophic for metabolites exported by other mutants), as well as a nine-species gut microbiome model.

## 1 Introduction

Interest in modeling microbial communities is growing as their impact on human health [11, 22, 23, 39] and the environment [16, 17, 30, 42] becomes more evident. The relationships between species in a community can show aspects of both cooperation, or dependence on the metabolites produced by other species, and competition for metabolites available in the environment. These interactions can lead to very different compositions of the microbial community, and understanding the underlying dynamics has important implications for potentially controlling the community composition by introducing new species or metabolites, or selectively eliminating certain species via antibiotics or immune action.

The simplest model for interaction between different species is the Lotka-Voltera model. This is a reduced-form model that summarises the interactions between species by an interaction matrix, and therefore is not able to represent interactions that are mediated by the exchange of metabolites [29]. Flux balance analysis (FBA) [32] and its many variants leverage genome-scale chemical reaction networks [43] to understand the interactions between microbial species at the level of metabolic interactions. FBA models typically assume that all species in the community cooperatively maximize a global community-wide objective function, possibly at the expense of lowering their own individual growth rates – notable exceptions being [6, 40] (see Section 1.1.2 for a detailed discussion). In this work, we propose a *non-cooperative* model where each species maximize its own grow rate, albeit taking into consideration the second-order effects of the metabolites it secretes into the environment. Our main contributions are as follows.

a. We propose a game-theory based model for predicting the composition of a microbial community. In this model, the various species are players in a non-cooperative game where the goal of the players is to maximize their own utility, which we define as their growth rate. However, unlike most game theoretic models, the “actions”, i.e. the reaction rates, feasible for a particular species depend on the actions taken by other species. We show that a generalized Nash equilibrium for this game exists, and can be efficiently computed by a globally convergent iterative algorithm that solves a convex quadratic program in each step. Our numerical experiments show that this algorithm scales to genome-scale metabolic models.
b. We consider the dynamic setting where biomasses of the species evolve according to the instantaneous growth rate given by the generalized Nash equilibrium of the game corresponding to the current biomass composition. Computing the steady state of these dynamics by solving the ordinary differential equations involves discretizing time and solving a Nash equilibrium problem at each time point – a very time-intensive procedure. We show that a steady state always exists, and that it can be efficiently computed by solving a single generalized Nash equilibrium problem. We also show how to test the robustness of a steady state to perturbations in the biomasses of each species, as well as to invasion by a new microbial species not currently in the community. We develop an algorithm to identify a steady state that is stable to biomass perturbation if one exists.
c. We discuss results of numerical experiments predicting the composition of a community of four *E. coli* mutants, with each mutant auxotrophic for a metabolite exported by one of the other mutants, and a nine-species gut microbiome model that can be directly compared to experimental data. We find that the steady state computed by a previously proposed community flux balance model that enforces cooperation is, in fact, unstable with respect to perturbations in the biomasses. We show that our approach implies that there are several qualitatively different steady states consistent with a given level of metabolite supply. This aligns well with studies showing that microbial communities can have a wide range of steady state behaviors even when the environmental conditions are the same [18, 33].

The rest of this paper is organized as follows. In Section 1.1 we review related literature and provide a background for our analysis. In Section 2.1 we introduce our game theory based model for predicting reaction rate profiles for each species in the community for a given biomass. In Section 2.2 we propose a model for the biomass evolution and show how to efficiently compute steady state biomass distributions. In Section 2.3 we characterize properties of stable steady states, and show how to use these properties to efficiently compute stable steady states. In Section 3 we use our procedure to predict biomass and reaction rate profiles for two model communities. Section 4 includes a discussion of our findings, a more in-depth analysis of how our model compares to two closely-related models [6, 8], and avenues for extending and improving our approach.

### 1.1 Related literature

Our work draws and builds on the following four different streams of literature.

#### 1.1.1 Flux balance analysis (FBA)

FBA is an approach for studying metabolic networks, i.e. the collection of metabolites found in an organism as well as the genes that encode enzymes that catalyze metabolic reactions. FBA uses genome-scale metabolic models and tools from optimization to compute the steady-state consumption and production of metabolites in an organism in order to predict the growth rate of the organism [32, 34]. A major benefit of FBA is that it only requires the stoichiometric constants of the metabolites in each reaction, rather than kinetic rate constants needed for a traditional dynamical systems approach to metabolic modeling that, in practice, are difficult to experimentally estimate [13]. The FBA approach has been gaining popularity in the last decade because high-quality metabolic network reconstructions for a number of different organisms have become available [24, 25, 31].

FBA imposes mass balance constraints that ensure the amount of each metabolite excreted and absorbed across all species are equal, as well as lower and upper bounds on the rate of each reaction. Reaction rates are calculated, subject to these constraints, so as to maximize an objective function. In single species models, this objective function is typically the rate of some “biomass” reaction that represents the consumption of metabolites needed for members of the species to reproduce. On the other hand, canonical multi-species FBA models are bi-level optimization problems wherein a community-level objective function is optimized in the outer problem, and each species maximizes its biomass reaction in the sub-problems [8, 47]. Our model is a departure from this last point – in our model each species maximizes its own biomass without any regard for any community level objective function, i.e. the species interact in a Nash equilibrium-like fashion. The importance of accounting for the behavior of selfish agents in community metabolism and biological systems in general has been increasingly recognized [6, 38], and requires game theoretical extensions of classic community FBA models.

#### 1.1.2 Microbial communities as a game

We model the steady-state composition of a community of microbial species as the equilibrium outcome of the competition between the various bacterial species for nutrients (i.e. metabolites). While not an FBA-based approach, Dubinkina et. al. [12] and Goyal, Dubinkina, and Maslov [19] predict the composition of the microbiome by proposing that the equilibrium composition is a stable matching between species and nutrients. Harcombe et. al. [20] consider competition between microbial species on a lattice, where space is the limiting resource and each species is described by an FBA model.

Combining FBA with game-theoretic/multi-agent system models has potential to be a useful approach to modeling the interaction between multiple bacterial species. Zomorrodi and Maranas [47] study the composition of a microbial community via a bi-level optimization model, referred to as OptCom, where each species maximizes its own growth rate while simultaneously cooperating with the other species to maximize the overall community growth rate. Our model can be considered an extension of OptCom that considers the Nash equilibrium behavior of the community compositions without consideration for a community-level objective function, accounts for and explicitly calculates the biomasses of each species in the community, and considers the dynamic stability of computed solutions. The solution of OptCom also entails solving a large bi-level non-convex problem, whereas our model only requires solving a sequence of single-level quadratic programs.

The SteadyCom model presented by Chan et. al. [8] is another modification of OptCom that specifically addresses the issues of simultaneously computing species biomasses and computing solutions in a more tractable manner. SteadyCom assumes that, at steady-state, each species has the same biomass-weighted reaction flux and that the different species cooperate to maximize their common biomass reaction rate. With these assumptions the bi-level problem reduces to solving a sequence of linear programs. Unlike our model, SteadyCom does not consider Nash equilibrium behavior from the species, nor does it consider dynamic stability of its solutions. The model in [40] is also a multi-level model that first identifies a species that can grow off the supplied nutrients and solves the corresponding FBA problem; next, it works with species that can grow off the metabolic by products of the first set of species, until the total community biomass is optimized.

Zomorrodi and Segre [48] used FBA to compute the payoff matrix for a producer-consumer game amongst *n* bacterial species. While Nash equilibria for this game are efficiently computed for moderate-size payoff matrices via an integer linear programming formulation, the size of the payoff matrix grows exponentially in the number of bacterial species as well as the number of metabolites considered. The model also focuses on the presence or absence of reactions producing metabolites of interest, rather than studying the values of fluxes for each reaction in the entire set of reactions.

NECom [6] considers a game theoretic model very similar to the one proposed in this paper when the biomasses are fixed. There are subtle modeling differences between the two approaches that have important methodological consequences, in particular, in our approach a generalized Nash equilibrium is guaranteed to exist, whereas this might not be the case in NECom. Furthermore, [6] does not discuss biomass evolution dynamics. See Section 2.1.2 for a more detailed discussion of the two approaches.

We model the interaction between the species as a non-cooperative game in which each species attempts to maximize its growth rate while being constrained by FBA constraints that can depend on the fluxes of other species. We assume that the reaction rates are given by generalized Nash equilibria (GNE). This approach has the advantage of not imposing a community-level objective function on the individual species, allowing us to study the more realistic case where the microbial community can operate sub-optimally at the community level [36]. We also discuss the stability of the associated GNEs to perturbation in biomasses and invasion by new species.

#### 1.1.3 Dynamic FBA

Typically, the quantity of interest when analyzing a microbial ecosystem is the relative proportion of each species. However, FBA does not provide this information, since it focuses on fluxes of the biomass reactions and not the biomasses. Dynamic FBA (DFBA) [26] models the dynamics of the biomasses by a Lotka-Volterra model with the growth rates given by the rates of the biomass reactions as calculated by FBA at that point in time. DFBA has been extended to model cooperative community dynamics [46], and a number of efficient implementations have been developed [5, 21]. We model the dynamics of microbial communities by assuming that the instantaneous growth rates for all species are given by a generalized Nash equilibrium for the FBA problem. However, we do not explicitly simulate the full dynamics. In fact, one of our main results is that one can compute steady states by solving for the generalized Nash equilibrium of a related game. This is more efficient that simulating the full dynamics, and similar in spirit to how SteadyCom calculates community biomass composition.

#### 1.1.4 Generalized Nash equilibria

The available actions, i.e. reaction rates, for any player (species) in our model are limited by the supply of metabolites that are secreted by or shared with other species. Consequently, the actions for each player depend on the actions of the other players. Nash equilibria for such games are called generalized Nash equilibria (GNE), and the associated games are called generalized Nash equilibrium problems (GNEP). We consider GNEPs where the objective and the constraints are described by linear functions. Several different approaches exists for solving such games: solving the concatenated KKT conditions for all players [10], formulating the GNEP as the minimization of a Nikaido-Isoda function [45], and penalty methods that solve a sequence of simpler Nash equilibrium problems [14]. We use a method that iteratively solves, in parallel, a quadratically regularized version of the best response problem of each player with the other players’ strategies fixed [15]. This approach is guaranteed to converge to a GNE, and we find in our numerical experiments that the method scales well as the number of species increases.

## 2 Methods

### 2.1 Model

We start by formulating a GNEP assuming that the biomass for each species is fixed. We consider the dynamics of the biomasses in Section 2.2.

#### 2.1.1 Chemical reactions

Let [*K*] = {1, 2,…, *K*}denote the set of *K* microbial species present in a compartment or community. Let *x* = [*x_k_*]_k∈K_ denote the biomasses of the *K* species. In this section, we assume that *x* is fixed. Let [*I_k_*] = {1,…, *I_k_*} index the set of metabolites in species *k* ∈ [*K*]. Let [*I_c_*] = {1,…, *I_c_*} index the set of metabolites found in the common space shared by the microbes^1^. In Figure 1, *K* = 2, *I*_1_ = 4, *I*_2_ = 3 and *I_c_* = 3.

**Figure 1:**
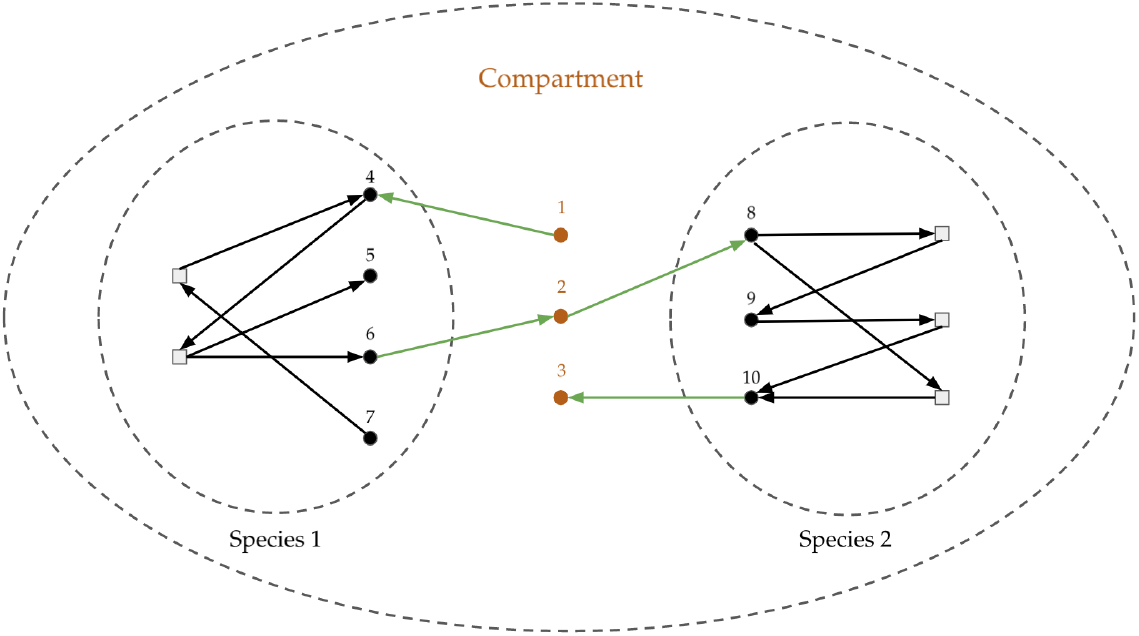
Schematic of microbial interactions, with grey squares representing reactions consuming and producing metabolites, represented by circles. Orange circles denote metabolites in the shared compartment, whereas black circles represent metabolites within a species. Arrows represent metabolites taking place in a reaction, with an arrow from (towards) a metabolite representing consumption (production) of that metabolite by the reaction. Green arrows represent exchange reactions that move metabolites between the shared compartment and a species, whereas black arrows represent reactions within a species. Note that the same type of metabolite is tracked as multiple different metabolites i.e. metabolite 2 in the shared compartment is the same type of metabolite as metabolite 6 in species 1 and metabolite 8 in species 2.

There are two sets of reactions associated with each species *k* ∈ [*K*]. The first set of reactions [*J_k_*] = {1,…, *J_k_*} change the concentration of metabolites within an individual cell of species *k*. The second set of reactions 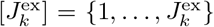 are exchange reactions, i.e. these ingest or excrete metabolites from the cell to the shared compartment. In Figure 1, black lines schematically denote the internal set of reactions [*J_k_*] and the green lines indicate the exchange reactions 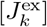.

Let 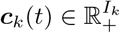 denote the metabolite concentrations in species *k* at time *t*. Then

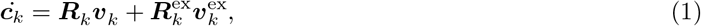

where 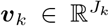 (resp. 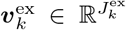) denotes the rates of the internal (resp. exchange) reactions, and the stochiometric matrices 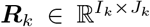 and 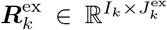 detail the impact of the reactions on the metabolite concentrations. The (*i,j*)-th element *R_k_*(*i,j*) of ***R**_k_* denotes the change in the concentration of metabolite *i* when reaction *j* proceeds at a unit rate: *R_k_*(*i,j*) > 0 if the metabolite *i* is produced by reaction *j* and *R_k_*(*i,j*) < 0 if metabolite i is consumed by reaction *j*. The stochiometric matrices 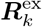 for the exchange reactions are special in the sense that every row has exactly one element equal to ±1, where *R_ex,k_*(*i,j*) = +1 (resp. *R_ex,k_*(*i,j*) = −1) indicates that the reaction *j* ingests (resp. excretes) metabolite *i* from the compartment, i.e. increases (resp. decreases) the concentration of the metabolite. At steady-state, each species must ensure that *ċ_k_* = 0 i.e.

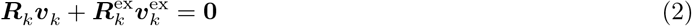

The fluxes within the species are further restricted by upper and lower bounds ***u**_k_* and ***ℓ**_k_*, respectively, that define maximum and minimum allowable fluxes, determined by the chemistry of the reactions

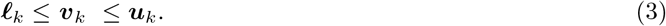

##### Assumption 1

(Bounded fluxes). *The bounds* (3) *and the flux balance constraints* (2) *imply that the set of feasible values* (*v_k_*, 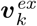) *are bounded for all k* ∈ [*K*].

The exchange reactions result in a change in concentration of the metabolites in the compartment. Let 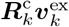, for 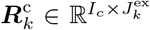, denote the metabolites ingested by each cell of species *k*. Note that 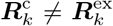 since the metabolites might be indexed differently in species *k* and the compartment. We assume that the inflow rate of metabolites into the compartment is fixed and is given by 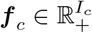. Thus, we require that

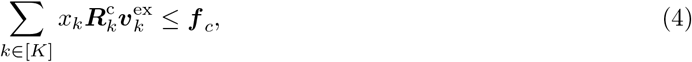

where *x_k_* denotes the biomass for species *k*. By allowing an inequality above, we are implicitly assuming that any unused metabolite diffuses away. The constraints (2)-(3) are species specific constraints, whereas (4) is a constraint *across* species.

#### 2.1.2 Nash equilibrium

The goal of the model is to predict the reaction rate profiles (*v*, *v*^ex^) that are likely to be observed in experiments. To this end, we posit that (*v*, *v*^ex^) corresponds to a Nash equilibrium of an appropriately defined game where the “players” are the species. This approach is similar to that in [6]. We will comment on the differences between the two approaches later in this section.

For given biomasses *x*, the set of feasible reaction rate profiles (*v*, *v*^ex^) for all species is given by the set

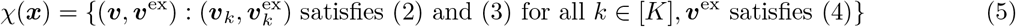

We assume that the payoff for species *k* is given by

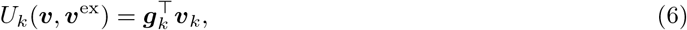

where ***g**_k_* denotes a combination of reactions that lead to biomass growth, and the set of feasible reaction rates for species *k* given the rates of all other species *ℓ* ≠ *k* is given by the set

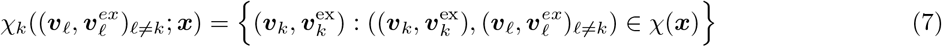

Let [*K*(*x*)] = {*k*: *x_k_* > 0} denote the set of species with positive biomasses. A reaction rate profile (*π*, *π^ex^*) is a GNE for the game defined by (6) and (7) if there is no incentive for any species *k* ∈ [*K*(*x*)] to deviate from (*π_k_*, 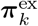), i.e. for all *k* ∈ *K*[*x*],

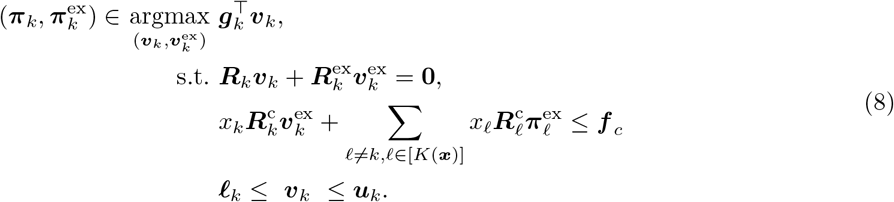

Our first result is that such a GNE exists.

##### Lemma 2.1

(Theorem 1 in [35]). *A GNE exists for the game* (8) *for any x and any chemical reaction parameters satisfying Assumption 1.*

We describe an algorithm for computing a GNE in Section A.1. Our game formulation is identical to the NECom algorithm proposed in [6], with one notable difference. In NECom, the constraints on the exchange reactions in the best response problem (8) are

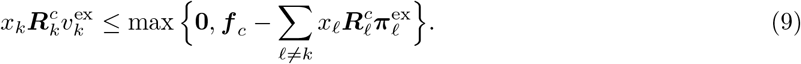

The rationale for introducing (9) is that it does not force altruism between the species i.e. no species is forced to produce metabolites for another species when doing so does not provide an immediate personal utility gain. However, introducing (9) results in serious methodological consequences. The best-response problem in NECom is non-convex, and therefore, Lemma 2.1 does not guarantee that a GNE exists. Furthermore, even if a GNE were to exist, the non-differentiability of (9) will make establishing stability of the GNE nearly impossible. It is also not clear whether imposing (9) is appropriate from a modeling perspective. Prohibiting forced altruism can cause the best-responses to be too myopic, and, as a consequence, rule out reaction profiles that are experimentally observed. For example, NECom when directly applied to auxotrophic communities, e.g. the four *E. coli* community analyzed in Section 3.1, predicts that the only possible community composition is the one where each species has *zero* biomass. In [6], this issue is handled in an ad hoc manner by introducing the additional objective of maximizing the total community biomass; it also employs a dFBA-like model with an NECom model solved at each time step and an initial supply of the auxotrophic metabolites. In contrast, our model is able to calculate non-zero steady states for this auxotrophic community without adding a community objective function, using dFBA, or having an external supply of the auxotrophic metabolites. Recall that our goal is to design a model that reliably predicts observed reaction rate profiles. In this context, the enforced altruism implicit in (8) is a better model since it is able reproduce observed behavior without any additional corrections.

### 2.2 Population dynamics and steady states

In this section, we describe a model for biomass evolution. These dynamics are similar to those in SteadyCom [8], but with the biomass fluxes calculated using the Nash equilibrium approach described in Section 2.1.2 i.e. we assume that at each point in time the vector of reaction rates is a GNE with respect to a biomass-weighted FBA game. Note that the model in this section is an NECom model where the biomasses are not fixed parameters, but rather evolve over time.

We assume that the quantity of each species of bacteria evolves according to the following Lotka-Volterra system of equations

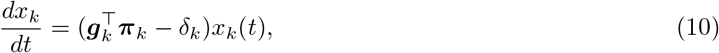

where *δ_k_* denotes the death rate for species *k* and 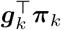 denotes the growth rate for species *k* corresponding to a GNE (*π*, *π*^ex^) of the game where the species biomasses are given by *x*(*t*). We are interested in efficiently computing steady states lim_t→∞_ *x*(*t*) and analyzing their stability. Although our focus is on the steady state behavior of (10), temporal dynamics of (10) can be easily computed by discretizing time, and computing a GNE for (8) at every time step.

Next, we show how to compute the steady states directly, i.e. without solving (10). We start with the following definition.

#### Definition 2.1.

(*x*, *π*, *π*^ex^) *is a steady state* GNE *if* (*π*, *π^ex^*) *is a GNE for the x-weighted game, and 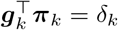 for all k* ∈ *K*[*x*].

The following result characterizes the set of steady states for (10).

#### Lemma 2.2.

*x is a steady state for* (10) *if, and only if, there exist* (*π*, *π^ex^*) *such that* (*x*, *π*, *π^ex^*) *is a steady state GNE as defined in Definition 2.1*.

Lemma 2.2 allows us to efficiently check whether a given *x* is a steady-state for the dynamics in (10). However, we do not yet know how to efficiently compute a steady-state *x*. Next, we introduce a game that allows us to identify candidates for the steady-state. The players in this new quantity-weighted game are still the species, and the actions for each player are a triple of quantities (*x_k_*, *v_k_*, 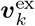), the utility of the player *k* is *U_k_*(*x*, *v*, *v*^ex^) = *x_k_*, and the set of feasible actions is

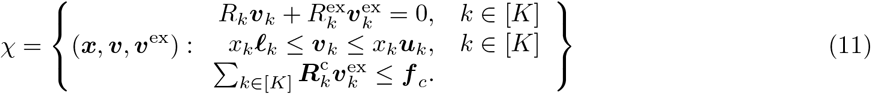

Thus, (*x*, *ω*, *ω*^ex^) is a GNE for this new quantity weighted game if, and only if,

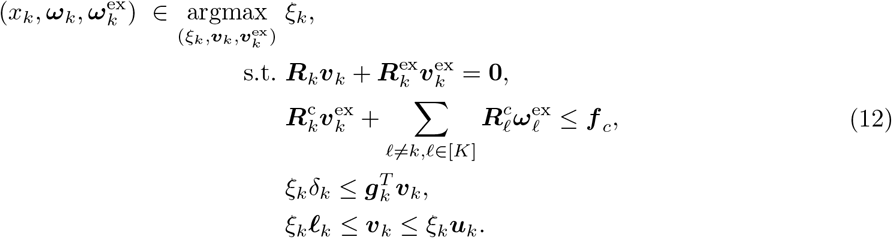

The next result establishes that a GNE (*x*, *ω*, *ω*^ex^) for (12) is a steady state GNE for (8), and therefore, *x* is a steady state for (10).

#### Lemma 2.3.

*Let* (***x***, ***ω***, ***ω**^ex^*) *be a GNE for the game* (12). *Define* 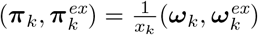 *for k* ∈ [*K*(*x*)]. *Then* (***x***, ***π***, ***π**^ex^*) *is a steady-state GNE.*

Since all constraints and objective functions for (12) are linear, a simple extension of Lemma 2.1 implies the following result.

#### Corollary 2.1.

*Suppose there exists* (***x***, ***v***, ***v**^ex^*) ∈ *χ with* 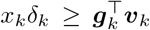 for all *k*. Then a GNE for the game (12) *exists*.

A GNE for (12) can be efficiently computed by solving a sequence of quadratic programs, see Section A.1 in the appendix for more details. Ideally, one would like the set of GNEs for (12) to coincide with the set of steady state GNEs for (8). This is not necessarily the case. However, we are able to establish that the set of GNEs of (12) is a super set of steady state GNEs that are *stable* with respect to biomass fluctuations.

### 2.3 Stability to perturbation in biomass and invasion

In Section 2.2 we showed how to compute one possible steady state for the Lotka-Voltera dynamics (10). However, this steady state may be unstable to fluctuation in the biomasses. Such an unstable steady state is unlikely to be experimentally observed, and therefore, we need a methodology for identifying stable steady states. In the next section, we first describe how to check stability of a given steady state, and then describe an algorithm that can identify if one exists. In Section 2.3.2 we show how to check whether a given steady state is stable under invasion by another species.

#### 2.3.1 Stability of steady state to fluctuations in biomass

Let ***x*** denote any steady state for 10. Then, there exists (***π***, ***π***^ex^) such that (***x***, ***π***, ***π***^ex^) is a steady-state GNE for (8). We want to understand the stability of this steady state for fluctuations in the biomass ***x***, since stability is a crucial property of microbial ecosystems [9]. Suppose the steady state ***x*** is perturbed to ***x***+***p***(0), where ***p***(0) ≈ **0**. We are interested in understanding the dynamics ***p***(*t*) for *t* ≥ 0. In particular, whether ***p***(*t*) → 0, and thereby keeps the community composition stable. Since the dynamics (10) are defined by Nash equilibria, we are effectively interested in understanding the comparative statics of the Nash equilibria [7].

We first discuss our approach informally and then state the formal mathematical result. Let 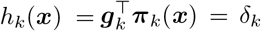 denote the growth rate at the steady state ***x***, where we emphasize that the GNE ***π*** is a function of ***x***. Then

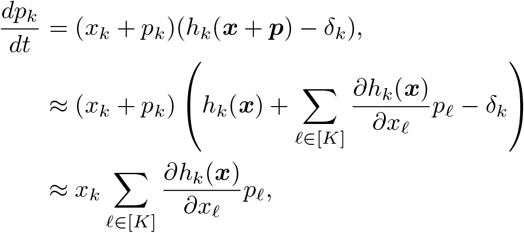

where we assume that the perturbation is small enough so that quadratic terms in ***p*** can be ignored, and use the fact that *x_k_*(*h_k_*(***x***) – *δ_k_*) = 0 for all *k*. Thus, we have that

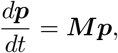

where 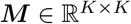 with the (*k,j*) element 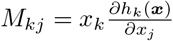.

##### Lemma 2.4

(Theorem 9.3.2 in [3]). *The steady state **x** is stable to fluctuations in the biomass if the real part of all the eigenvalues of **M** are all less than* 0.

In Lemma A.1 we show how to compute 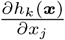 for all *k*, *j*, in order to define **M**.

Next, we establish a partial converse for Lemma 2.3. As a first step, we define a subset of steady state GNEs.

##### Definition 2.2

(Resource-constrained steady-state GNE). *A steady-state GNE* (***x***, ***π***, ***π**^ex^*) *is* resource-constrained *if for all species k, the optimal value of the best response problem (8), when the biomasses x_ℓ_ and fluxes 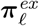 for all species ℓ* ≠ *k are fixed, and the biomass of species k is set to z_k_* > *x_k_, is less than δ_k_*.

Lemma A.2 establishes that at a resource-constrained GNE each species’ growth is limited by the consumption of at least one external metabolite. However, this is not a sufficient condition for a GNE to be resource-constrained. For that, we must have that increasing the biomass to *z_k_* > *x_k_* forces a decrease in the uptake of a metabolite that ultimately results in a decrease in the rate of biomass producing reactions. A similar concept was formalized for the computation of steady states in SteadyCom (see the proofs of Theorems 1 and 2 in the Appendix of [8]).

##### Lemma 2.5.

(***x***, ***π***, ***π**^ex^*) *is a resource-constrained steady-state GNE if and only if* 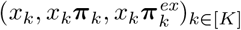 is a GNE for (12).

Thus, we have a partial converse for Lemma 2.3. Next, we show that stable steady state GNEs are a subset of resource-constrained steady states, and therefore, a subset of GNEs for (12).

##### Lemma 2.6.

*Suppose* (***x***, ***π***, ***π**_ex_*) *is a steady state GNE, but it is not resource-constrained. Then it is unstable to perturbation in the biomass **x***.

Since observing a microbial community at an unstable equilibrium is unlikely, Lemma (2.6) allows us to focus our attention on resource-constrained steady state equilibria, and therefore, justifies using (12) to compute steady states. Recall that Corollary 2.1 establishes that the game (12) has a GNE. However, there are no general results that guarantee the existence of a stable steady state. In Appendix A.4 we develop an algorithm that exhaustively lists all possible candidates for steady state GNEs.

##### Lemma 2.7.

*There exists an exhaustive search algorithm to check for the existence of stable steady state GNE that terminates in finite time.*

The search algorithm is not scalable to very large communities, since it involve enumerating the vertices of high-dimensional polytopes. However, it is highly parallelizable, and therefore, the existence of stable steady state GNE can be verified given sufficient computational resources. See Appendix A.4 for more details.

#### 2.3.2 Stability to invasion

In the previous section, we were concerned with stability with respect to fluctuations in the biomass ***x***. Here, we consider stability of a steady state to invasion by a new species. Let (***x***, ***π***, ***π***^ex^) be a steady-state GNE for the *K* species, and index this new species by *K* + 1. Let *h*_*K*+1_(***x***, *ϵ*) denote the growth rate of the invading species when its biomass *x*_*K*+1_ = *ϵ*. Then species *K* + 1 successfully invades if lim_*ϵ*↓0_ *h*_*K*+1_(***x***, *ϵ*) > *δ*_*K*+1_. Lemma A.4 establishes that lim_*ϵ*↓0_ *h*_*K*+1_(***x***, *ϵ*) can be computed by solving an optimization problem.

## 3 Results

In this section, we use our proposed methodology for computing the steady state biomass and reaction profiles for two different bacterial communities: the four auxotrophic modified *E. coli* species and the nine species gut microbiome model studied in [8].

The results in Section 2.2 establish that a steady state for (10) can be computed by solving for a GNE of the game (12), and Lemma 2.4 allows us to check whether the steady state is stable. Furthermore, the exhaustive algorithm in Appendix A.4 is able to search for all possible stable steady states, if any, for (10). However, the running time of this algorithm was prohibitive even for the 4 species community. Instead, we used a sampling based algorithm to identify steady states. We sample *N* biomass vectors {***x***^(*s*)^: *s* = 1,…, *N*} and then call Algorithm 1 on each of the samples. We found that this procedure efficiently identified a stable steady state with *N* = 2^11^ for the *E. coli* model and *N* = 100 for the nine species gut model. Therefore, we did not have to use the exhaustive search algorithm described in Appendix A.4.

### Algorithm 1 GameCom

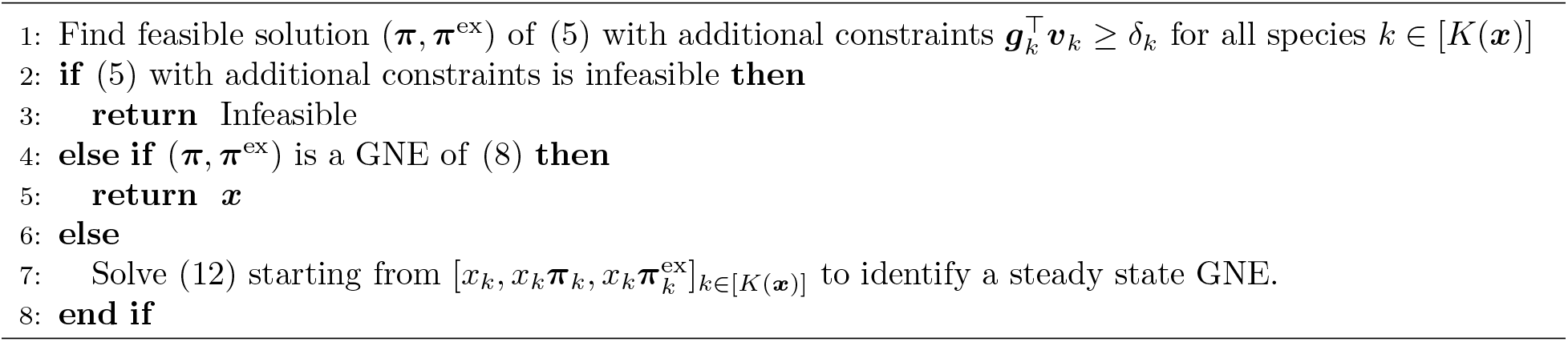

SteadyCom [8] and NECom [6] are two other algorithms available in the literature to compute reaction rate profiles. In contrast to our approach, SteadyCom assumes that all the species collectively maximize their common growth rate. NECom computes reaction rate profiles by solving for a GNE for given biomasses ***x***; however, with a different best-response optimization – see the discussion below (8) in Section 2.1.2. The fluxes computed by NECom for the 4 bacteria *E. coli* community are different enough from our results that we have ommitted a direct comparison to NECom in the results section. We comment on the differences between NECom and GameCom in Section 4.

### 3.1 Auxotrophic *E. coli* community

The community was created by taking four copies of *E. coli* metabolism and inhibiting the export and production of specific metabolites in each copy so that each of the four modified models can only grow in the presence of the other three. More specifically:

1. EC1 is able to synthesize arginine and phenylalanine and export arginine.
2. EC2 is able to synthesize lysine and methionine and export lysine.
3. EC3 is able to synthesize lysine and methionine and export methionine.
4. EC4 is able to synthesize arginine and phenylalanine and export phenylalanine.

Note that, in spite of the interdependence, there is also a degree of competition, e.g. EC1 and EC4 compete for lysine and methionine, and all four species compete for common resources, e.g. glucose. Metabolites necessary for the community to grow are externally supplied; we use the “western diet” provided in [8]. SteadyCom computes a steady-state for this model with a common growth rate of 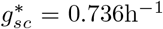. In this steady-state, the proportion of EC1 is 25.3%, EC2 is 32.5%, EC3 is 18.5%, and EC4 is 23.7%.

#### 3.1.1 Analysis of SteadyCom steady state SC

The first numerical experiment was to check whether the SteadyCom steady state (referred to as “SC”) is a steady state in our computational framework, and if so, whether it is stable.

SteadyCom assumes that the death rate for each species is identical, and predicts the proportion of each species; therefore, the death rate is not required for the calculations. We predict the absolute biomasses and account for the slow down in growth as the biomasses increase but the nutrient supply remains constant. Consequently, we need to explicitly specify the death rate. In order to compare our predictions, we also assume that the death rate of each of the species is identically equal to *δ*. In order to ensure that there is a non-zero steady state, we ensure that 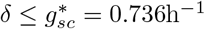, the maximum SteadyCom growth rate. We found that our results were robust to the specific value of *δ*. For our numerical experiments we set *δ* = 0.5h^-1^, and then rescaled the SteadyCom biomass proportions and reaction fluxes to obtain a set of biomasses and fluxes that are feasible for (8). We find that this re-scaled SC is a steady-state GNE, but it is *unstable* to perturbations in the biomasses.

#### 3.1.2 Steady states robust to perturbation in biomass *x*

Next, we compute other steady state GNEs for the problem and analyze their stability. We sampled *N* = 2^11^ = 2048 biomasses for the four species uniformly from [0,1]^4^ using a Sobol sequence generator [41]. Of these, 408 samples remain after Step 2 in Algorithm 1. Each of these samples yielded a steady state GNE either in Step 4 or Step 7 of Algorithm 1. The screening step on all of the *N* = 2^11^ biomass samples took approximately 1.9s per sample. Computing the steady state GNE for each of these 408 samples took approximately 3.5s on average, and checking for stability took 8.5s on average.

##### Very few stable steady states

Our first result is that, although there are many steady state GNEs, very few of them are actually stable – only 4 of the 408 steady state GNE are stable to biomass perturbations. Thus, stability is a very strong selection criterion, and the analysis of the results of the model reduces to only a handful of steady states. In the left panel of Figure (2)(a), we compare the total community biomasses for stable and unstable steady states. We find the range for the total biomass is much larger for the unstable steady states when compared to stable steady states. As expected, the steady state SC has the largest community biomass (see vertical dotted line in the left-hand plot).

**Figure 2:**
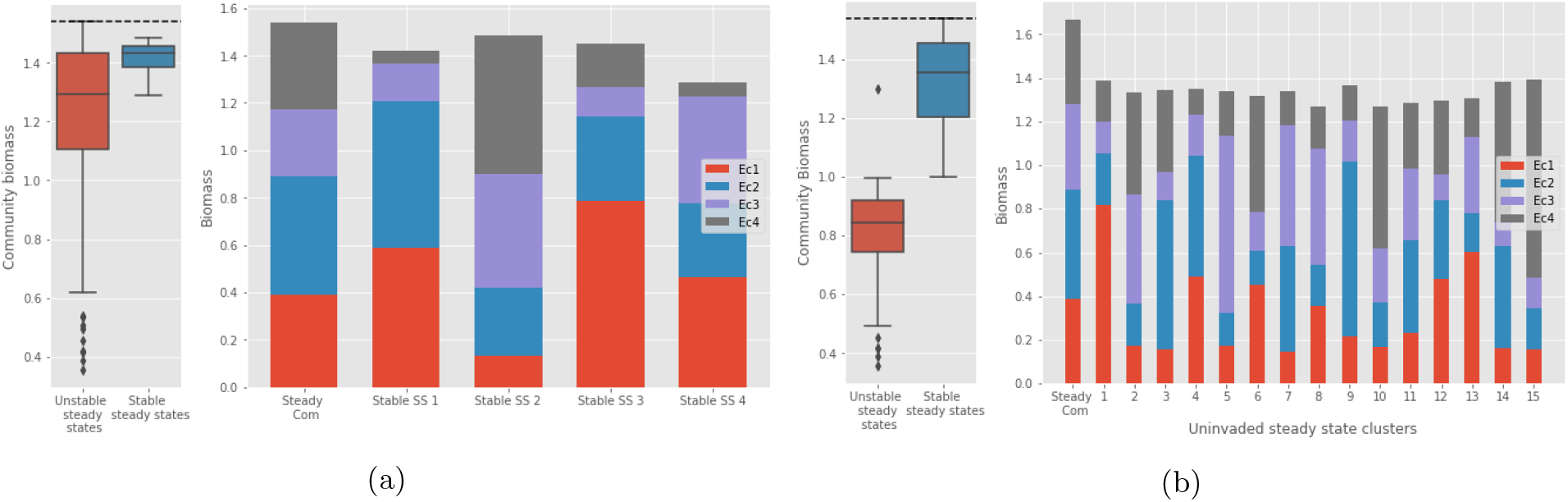
(a) Results for the community of four *E. coli* mutants. The box plots on the left the total community biomass for the four mutants. The bar charts on the right compare four stable steady states with the (unstable) SteadyCom steady state SC. (b) Compares steady states stable to invasion. The bar charts on the right compare steady states stable/unstable to invasion. The 333 steady states stable to invasion are grouped into 15 clusters.

In the right panel in Figure (2) (a) we plot the (unstable) steady state SC, and the 4 stable steady states computed by GameCom. We see that the 4 steady states are very different from each other, and also from SC. Thus, we find that assuming perfect cooperation, as in SteadyCom, does not predict the range of behavior that is possible in the microbial community.

##### Biomass distribution across stable steady states

Next, we carefully examine the variation in the biomass distributions for SC, SS2 and SS3. The goal here is to understand how two very distinct steady states can both be stable, and how they compare to SC. In Table 1 we list the externally supplied metabolites that are limiting, i.e. the corresponding compartment constraint is tight, and also the number of reactions that are equal to their upper or lower bounds, i.e. the bounds are active. We find that in all three steady states the four metabolites that are part of the auxotrophic interdependence between the species are fully utilized, and, among the externally supplied metabolites, only glucose is fully utilized. We also find that many more of the reaction bounds are active in SS2 and SS3, as compared to SC. Furthermore, we find that the very different steady states can be supported because even a small amount of key metabolites (e.g. arginine EC1, or lysine EC2, etc.) can sustain a relatively large quantity of the other species dependent on the key exported metabolite. Therefore, while the existence of each species is essential for the existence of the other species, they are not severely growth limiting, allowing for qualitatively different steady state GNE.

**Table 1:** Comparison of metabolite utilization and internal reaction fluxes for the second stable steady state in the right-hand part of figure (2), the third stable steady state from figure (2), and the SteadyCom steady state. The first column lists which of the four metabolites that the species are auxotrophic for are fully utilized i.e. 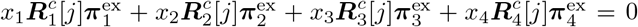, *j* being corresponding to the fully utilized metabolite. The second column lists the externally supplied metabolites that are fully utilized i.e. metabolites *j* where ***f**_j_* > 0. The third and fourth columns list, for each species, the number internal reactions that are at their lower bound and upper bounds, respectively.

|  | Limited auxotrophy metabolites | Limited external resources | Number of active lower bounds | Number of active upper bounds |
| --- | --- | --- | --- | --- |
| Stable steady state 2 | Arg, Met, Lys, Phe | Glucose | Ec1: 12, Ec2: 12<br>Ec3: 11, Ec4: 15 | Ec1: 10, Ec2: 12<br>Ec3: 11, Ec4: 15 |
| Stable steady state 3 | Arg, Met, Lys, Phe | Glucose | Ec1: 12, Ec2: 13<br>Ec3: 12, Ec4: 15 | Ec1: 8, Ec2: 13<br>Ec3: 14, Ec4: 17 |
| SteadyCom steady state | Arg, Met, Lys, Phe | Glucose | Ec1: 0, Ec2: 2<br>Ec3: 2, Ec4: 0 | Ec1: 0, Ec2: 3<br>Ec3: 3, Ec4: 1 |

##### Instability of SC

To understand why SC is unstable, we compare its fluxes to those in SS3. As noted earlier, in both SC and SS3, the four auxotrophic metabolites and glucose are fully utilized. Therefore, no species can simultaneously increase the rate at which it takes up one of these metabolites, and keep its biomass the same. As a rule of thumb, this will prohibit perturbations with increases in the biomasses to drive the biomass away from the steady state. In contrast to SC a number of the SS3 fluxes are at their upper or lower bound. For each *E. coli* mutant, a handful of these reactions are involved in carbon metabolism. For example, in EC1 the rate for the reaction catalyzed by glycogen branching enzyme is at its maximum possible value; consequently, in EC1 the rate of conversion of glucose to glycogen is limited even if more glucose becomes available. This prevents perturbations where species biomasses are decreased from destabilizing the steady state. As a simple example of this phenomenon, suppose the biomass of EC1 at SS3 is decreased while the biomasses of the other three species are held constant. The decrease in the biomass of EC1 results in more carbon available for the other three species; however, these species cannot utilize the available carbon because the rate of some reaction involved in glucose conversion is at its upper bound. The unused carbon is completely available for EC1 to grow back to its original biomass level. None of the carbon metabolic pathways are at their maximum fluxes in SC; therefore, such a perturbation can potentially be unstable. In a sense, SC is unstable because it is too efficient – in maximizing the community growth rate, the species’ reactions had to be perfectly balanced, and as a result the community is capable of utilizing additional resources that become available due to perturbations.

#### 3.1.3 Steady states stable to invasion

Next, we consider stability to invasion by a new species for each of the 408 steady states. In particular, we explore whether an *E. coli* mutant that can produce arginine, methionine, lysine, and phenylalanine, but not export any of these four metabolites, can invade. Computing the stability for each of the 408 steady states took approximately 25s per steady state. In contrast to stability to perturbations in the biomasses, stability to invasion by this fifth *E. coli* species is much more common - 333 of the 408 steady state GNEs cannot be invaded by new mutant. Note that the four stable steady states found in the previous section, as well as SC, are all stable to invasion. The results in this section are summarized in Figure 2 (b). On the left panel, we plot the biomass distribution for steady states that are stable/unstable to invasion. On the right panel we plot the steady states resistant to invasion clustered into 15 groups. As was the case with stability to biomass perturbations, steady states resistant to invasion have a larger community biomass. In contrast to stability to biomass perturbations, the stability to invasion is determined by the availability of externally supplied metabolites: stable steady states have at least one fully exhausted externally supplied metabolite whereas unstable steady states do not. The exhausted nutrient prevents the growth of the invading species.

Figure (3) plots the results for the death rate *δ* = 0.4h^-1^. We note that the results are qualitatively similar to those in Figure 2 that corresponds to a death rate *δ* = 0.50h^-1^, in the sense that there are multiple stable steady states with a large variation in the biomass distribution. There are more steady states when the death rate is lower: 753 steady states, 7 of which are stable to perturbations in the biomass ***x***. This increase is in line with the flux variability analysis summarized in Figure 2 of [8]. Increasing the death rate to 0.60h^-1^ resulted in only 120 steady states, and none of them were stable. Hence, establishing that it is more difficult to sustain all four species as the death rate increases.

**Figure 3:**
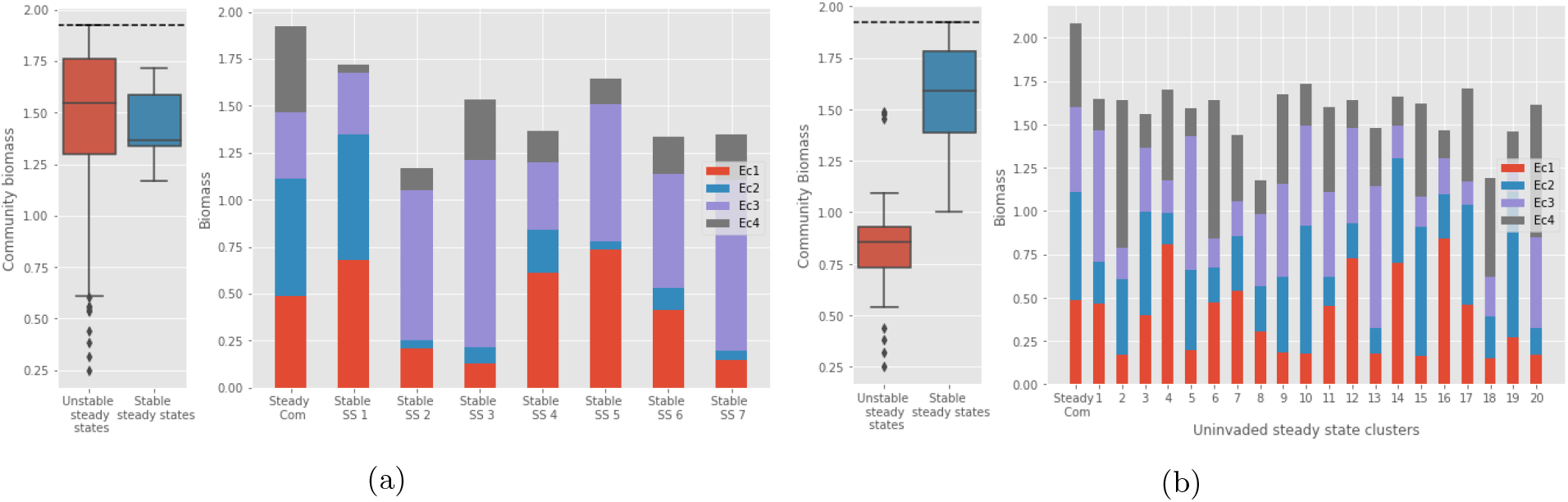
Results for death rate *δ* = 0.40 h^-1^. See footnote for Figure 2 for explanation of the plots.

### 3.2 Nine species gut microbiome model

Next, we apply our procedure to identify stable steady states for the gut microbiome model analyzed in SteadyCom [8]. The model consists of nine species comprising four phyla, as described in Table 2. We use the same model files for each species as used in SteadyCom, and use the same “Western diet” as the auxotrophic *E. coli* community in the previous section. In addition to being a more realistic community, the results of this model can be directly compared to experimental data from [28, 44].

**Table 2:** Species used in gut microbiome model

| Species | Phyla |
| --- | --- |
| B. theta | Bacteroidete |
| E. rectale | Firmicute |
| F. prausnitzii | Firmicute |
| E. faecalis | Firmicute |
| L. casei | Firmicute |
| S. thermophilus | Firmicute |
| B. adolescentis | Actinobacteria |
| E. coli | Proteobacteria |
| K. pneumoniae | Proteobacteria |

We sampled *N* = 100 biomass vectors close to the experimentally observed data in [28, 44] as candidate steady states. The biomass *x_k_* for each species *k* was chosen uniformly from an interval of width 0.1 centered at the average value for that species’ proportion in [28] and [44], with a minimum value of 0.001. Even though the values reported in [28, 44] are proportions, we treated them as absolute biomass weights. This is justified as long as the death rate is set close to the maximum growth rate computed in SteadyCom, since SteadyCom normalizes the biomasses to 1 and finds a growth rate that’s compatible with the absolute biomass values adding up to 1. The analysis in [8] shows a maximum possible community growth rate for this model is approximately 0.08h^-1^, therefore, we set the death rate to *δ* = 0.06h^-1^ for each species. From these 100 sampled biomass distributions, we were able to compute 20 steady states, and found that 2 of them were stable. See Figure 4 for the plots of the proportions of the different phyla for the two stable steady states. Also plotted are the phyla proportions as computed by SteadyCom and experimentally determined proportions from [28, 44].

**Figure 4:**
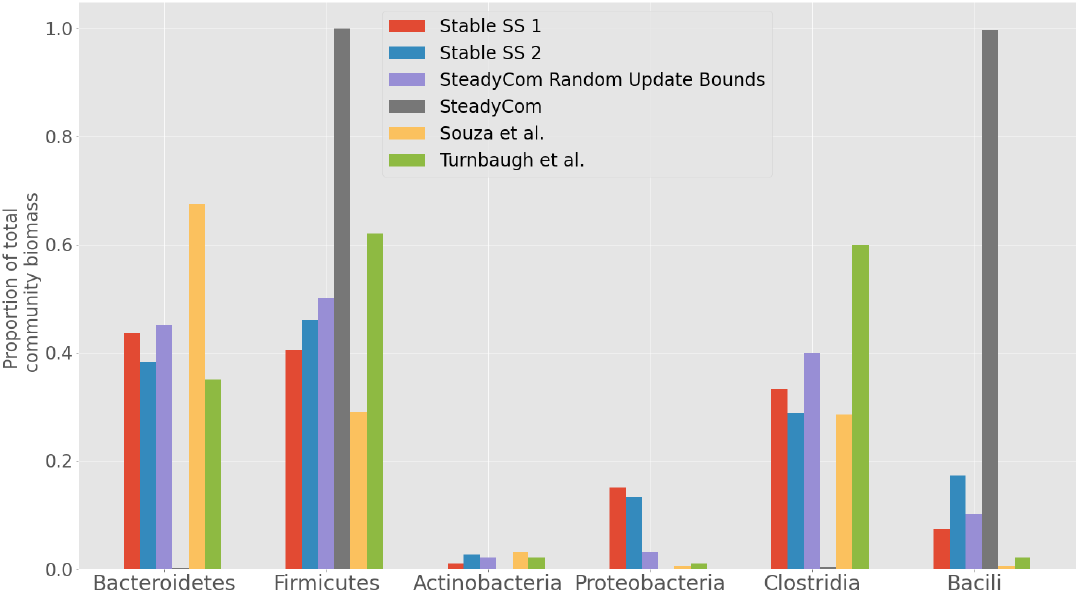
Stable steady state phyla proportions computed by our method (red and blue bar plots). Phyla proportions computed using SteadyCom without modifications to the model (grey bars) and with restrictions on the maximum fiber uptake rate (purple bars). Experimental data from [28] (yellow bars) and [44] (green bars) included for comparison. Two classes of firmicutes, clostridia and bacili, are also plotted since the majority of the firmicute population should be comprised of clostridia.

The predictions from our model match up well with the experimental results in [28, 44], with the exceptions of having too much proteobacteria (10% to 15% in our model compared to 1% to 2% in experiments), and the firmicutes being comprised of a larger fraction of bacili compared to clostridia.

Note that we did note take any additional ad hoc steps to improve the match between our results and the experimental data. SteadyCom requires ad hoc manipulations of the model to get similar results. The grey bars in Figure 4 plot the proportions computed by SteadyCom using the model and diet as is – the entire community is comprised almost entirely of firmicutes. This is because firmicutes grow most efficiently off of fiber, the main carbon source in the diet. SteadyCom is able to overcome this issue in one of two ways: by requiring that the proportions of all species are constrained to be sufficiently above zero, and/or imposing upper bounds on the rate of fiber uptake for each species. However, since there are no experimental measurements for these species-specific fiber uptake upper bounds, SteadyCom resorts to sampling many different fiber uptake bounds, and computes average species composition across the samples. The results of this are the purple bars in Figure 4 – putting a cap on the amount of fiber that any given species can uptake allows for multiple species to uptake the fiber needed for their growth. In contrast, our competition-based approach is able to obtain reasonable community proportions without artificially constraining uptake reactions with unknown upper bounds.

## 4 Discussion

We propose a new FBA model for predicting the composition of microbial communities. In our model the species in the community optimize their individual growth rate by competing for available metabolites. Therefore, we avoid imposing a community-level objective function that all species attempt to optimize in a cooperative manner. We show that a Generalized Nash Equilibrium (GNE) for the non-cooperative game always exists and can be computed by a globally convergent iterative algorithm that solves a quadratic program in each iteration. Depending on the chosen community-level objective function, computing a GNE can be significantly more efficient than solving the bi-level optimization problems encountered while computing the solution for a community level objective. SteadyCom simplifies to solving a small number of linear programs when the death rate for all species in the community is identical; however, when the death rates are species dependent one has to solve a non-convex bi-level optimization problem with no convergence guarantees.

Stability of a steady state to perturbations in the biomass composition and invasion from other microbial species is an important consideration for predicting community composition. This is because unstable steady states are unlikely to survive in biological environments that are inherently noisy. We propose a methodology for checking the stability of such steady states to both perturbation in biomasses and the introduction of new species. We also present an algorithm that can exhaustively list all possible candidates for steady states stable to biomass perturbation, and therefore, one can check whether a community has a such a stable steady state.

The numerical results reported in Section 3 confirm that removing the requirement that the members of the community must cooperate to optimize a shared objective allows a richer variety of steady state communities. This aligns well with studies showing that microbial communities can have a wide range of steady state behaviors even when the environmental conditions are the same [18, 33]. In our numerical experiments, we see that stability is a very strong selection criterion – most steady states are unstable.

GameCom shares a number of similarities to NECom, which have already been discussed in Sections 1.1.2 and 2.1.2. One of the differences between GameCom and NECom is that the shared metabolite constraints in NECom do not force the export of metabolites needed by other members of the community. Prohibiting forced altruism can cause the best-responses to be too myopic, and, as a consequence, rule out reaction profiles that are experimentally observed. For example, NECom when directly applied to auxotrophic communities, e.g. the four *E. coli* community analyzed in Section 3.1, predicts that the only possible community composition is the one where each species has *zero* biomass. In [6], this issue is handled in an ad hoc manner by introducing an additional objective of maximizing total community biomass; and employing a dFBA-like model with a NECom model solved at each time step and an initial supply of the auxotrophic metabolites. In contrast, our model is able to calculate non-zero steady states for this auxotrophic community without adding a community objective function, using dFBA, or having an external supply of the auxotrophic metabolites. Recall that our goal is to design a model that reliably predicts observed reaction rate profiles. In this context, the enforced altruism implicit in (8) is a better model since it is able reproduce observed behavior without any additional corrections.

While we are able to identify the set of steady states to biomass perturbations, we are not able to efficiently identify the particular steady state that is likely to be the convergence point for a given initial state, other than simply solving the ODE (10). We are able to predict whether a particular steady state is stable or unstable to invasion. However, we are not able to list all possible candidates for steady states that are immune to invasion, nor are we able to efficiently predict the new steady state, or whether or not the invading species survives. Understanding how to control the composition of a microbial community, by introducing new species or metabolites, changing the supply of metabolites, or changing the death rates, is a larger question that is both theoretically interesting and practically important. Analyzing the impact of the death rates in particular could be an interesting way of understanding the interaction between the microbiome and the immune system (e.g. intestinal toll-like receptors). We believe GameCom is a first step towards answering these questions.

## A Supplementary Materials

In this section, we include proofs for results presented in the main body of the paper, as well as additional results not included in the main body.

### A.1 Algorithm for computing GNE

In this section, we describe an algorithm for computing a GNE for the game (8); computing GNE for (12) is similar.

Define the operator ***T***(***v***, ***v***^ex^) = [*T_k_*(***v***, ***v***^ex^)]_*k*∈[*K*]_ as follows:

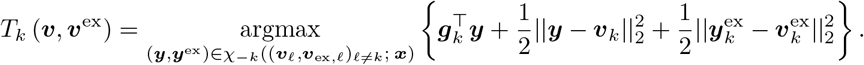

Note that the optimization problem implicit in the definition of *T* has a strongly convex quadratic objective function and linear constraints. Optimization problems of this form can be solved very efficiently [4].

Proposition 12.5 in [15] establishes that (***v***, ***v***^ex^) is a GNE if, and only if, (***v***, ***v***^ex^) is a fixed point of the operator ***T***. Proposition 12.17 in [15] applied to the problem defined here shows that ***T*** is non-expansive and, therefore, a fixed point of ***T*** can be computing using an averaging scheme of the form

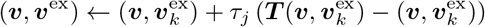

for a step length *τ_j_* ↘ 0 such that 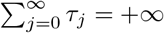.

In practice, using such an averaging scheme to find a GNE is slow because the regularization term and the step length *τ_j_* limit the change in (*v*, *v*^ex^) at each iteration to be small. To counteract this, we define a new operator 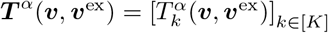 where

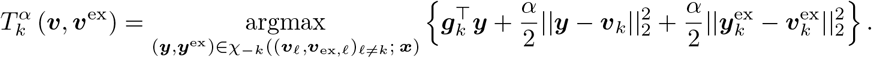

We compute a GNE by performing the averaging scheme with ***T**^α^*(***v***, *v*^ex^) where *α* = *α_j_* in iteration *j*, and it is a non-decreasing function of *j* that saturates at 1. By setting *α_j_* small for *j* small, we allow the reaction rates to take large steps, and hence, speed up the convergence to a fixed point. By ensuring that *α_j_* saturates at 1, we are guaranteed that the iterates converge to a fixed point of ***T***. Empirically, we found that we got very good performance by setting 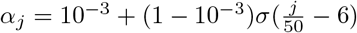, where *σ* denotes the sigmoid function.

### A.2 Calculating steady states

#### Lemma 2.2.

***x** is a steady state for* (10) *if, and only if, there exist* (***π***, ***π**^ex^*) *such that* (***x***, ***π***, ***π**^ex^*) *is a steady state GNE as defined in Definition 2.1*.

*Proof.* Suppose ***x*** is a steady state for the dynamics in (10). Then, for each *k*, either *x_k_* = 0, or *x_k_* > 0 and the growth rate of the species is equal to the death rate *δ_k_*. By definition, the growth rate for a species *k* is given by 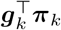 for some GNE equilibrium (***π***, ***π***^ex^) for an *x*-weighted game. Thus, it follows (***x***, ***π***, ***π***^ex^) is a steady state GNE.

Suppose (***x***, ***π***, ***π***^ex^) is a steady state GNE. Then we have that (***π***, ***π***^ex^) is a GNE for the *x*-weighted game, and 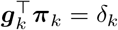. Thus, it follows that **x** is a steady state for the dynamics in (10).

#### Lemma 2.3.

*Let* (***x***, ***ω***, ***ω***^ex^) *be a GNE for the game* (12). *Define* 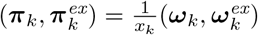 *for k* ∈ [*K*(*x*)].

*Then* (***x***, ***π***, ***π**^ex^*) *is a steady-state GNE*.

*Proof.* (***π***, **π**_ex_) is clearly feasible for (8) with weights *x*. Suppose however that it is not a steady-state GNE i.e. for some *k* there exists (*ϵ_k_*, 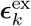) feasible for (8) such that 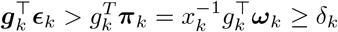.

Define 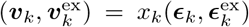, and 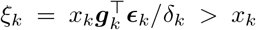. Since *ℓ_k_* ≤ 0 and *u_k_* ≥ 0, it follows that (*ξ*, *v_k_*, 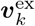) is feasible for (12). We have then that (*x_k_*, ***ω**_k_*, ***ω***_ex,*k*_) is not optimal for (12). A contradiction.

### A.3 Stability to perturbation in biomass

We need some notation in order to explain the construction of the system of equations for solving for 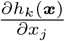.

i. Let

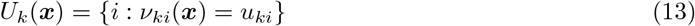

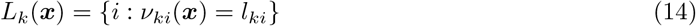

denote the indices for which the optimal solution ***π**_k_*(***x***) for the linear program (8) at biomasses ***x*** is equal to the reaction upper or lower bound. Let

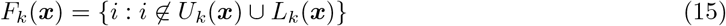

be the sets of free non-exchange reactions at the optimal solution for (8) at biomasses **x**.
ii. Let 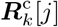 denote the *j*-th row of the matrix 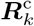. Let

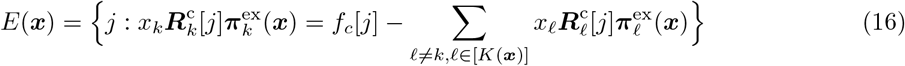

denote the set of community exchange constraints that are active in (8). Note that the set of active exchange constraints is independent of species *k* ∈ *K* [*x*].

The following characterizes the values of 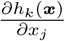 needed to construct **M**.

#### Lemma A.1.

*Suppose* (***x***, ***π***, ***π**^ex^*) *denotes any steady state GNE. Suppose the dual linear program of* (8) *has a unique solution for all k* ∈ [*K*]. *In particular, let **λ**_*k*_ be the dual optimal solution corresponding to the metabolite exchange constraints. Also define* 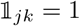 *if j* =*k*, 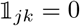 *otherwise. Then we have*

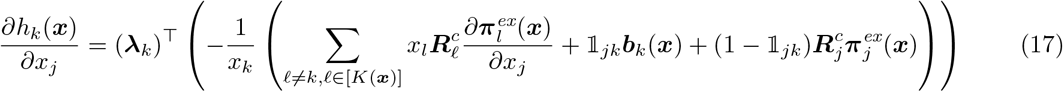

*where the partial derivatives of the fluxes with respect to changes in the biomasses are determined by solving the following system of equations over all k, j* ∈ [*K* (*x*)]

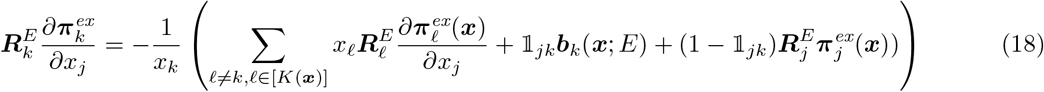

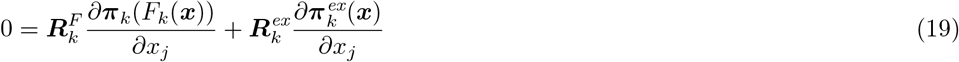

*Proof.* Recall the “utility” *h_k_*(*x*) for species *k* is given by the optimal value of the LP (8) at a steady state Nash equilibrium (***π***(***x***), ***π***^ex^(***x***)) with the biomasses set to ***x***. Let

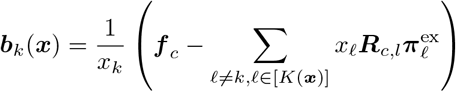

denote the right-hand side of the metabolite exchange constraints for the problem corresponding to species *k*. Since the dual solution is assumed to be unique, we have for all ***b**_k_* sufficiently close to ***b**_k_*(*x*) that the optimal values *h_k_*(***b**_k_*) = *h_k_*(***b**_k_*(*x*)) + (***λ**_k_*)^T^(**b**_k_ – ***b**_k_*(***x***)). Therefore,

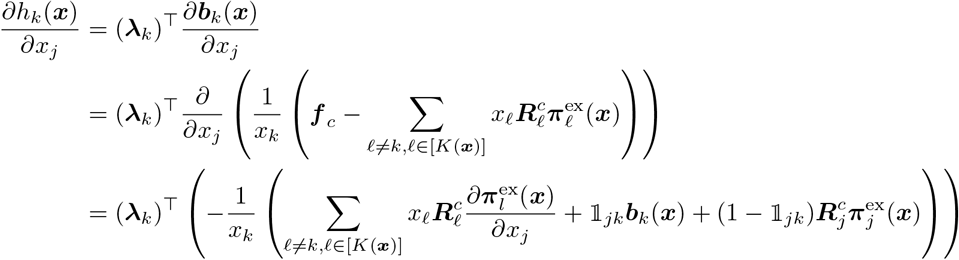

Thus, it follows that the optimal solution for ***b**_k_* will have ***π**_k_*[*U_k_*] = ***u**_k_* [*U_k_*(***x***)], ***π**_k_*[*L_k_*(***x***)] = ***ℓ**_k_*[*L_k_*(***x***)], and

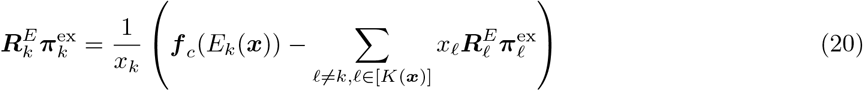

and the free variables are determined by the systems of equations

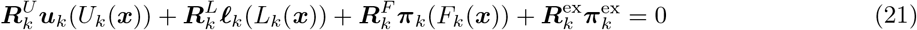

We can differentiate (20) and (21) to solve for 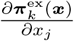 and 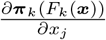 for all *k*, *j* ∈ [*K*(*x*)].

See Section A.5 for a discussion of the assumption that there exists a unique solution to the dual LP.

### A.4 Existence of stable steady state equilibria

The following result gives more context to the resource-constrained steady state GNE defined in Section 2.3.1.

#### Lemma A.2.

*Let* (***x***, ***π***, ***ν**^ex^*) *be a resource-constrained steady-state GNE. Suppose dual optimal solutions for each species’ best response problem (8) at* (***x***, ***π***, ***π***_ex_) *is unique. Let U_k_*(***x***), *L_k_*(***x***), *and E*(***x***) *denote the active sets defined in* (13)-(16). *Then for each species k* ∈ [*K*(***x***)], *there exists an active constraint j* ∈ *E*(***x***) *such that*

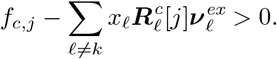

*Proof.* By contradiction suppose there exists a species *k* such that for all *j* ∈ *E*(***x***), we have that

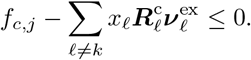

Recall the compartment exchange constraints imply that

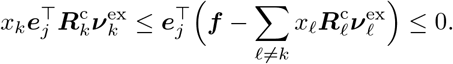

Since *x_k_* > 0, it follows that 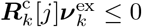 for all *j* ∈ *E*(*x*). Thus, for all *z_k_* ≥ *x_k_*

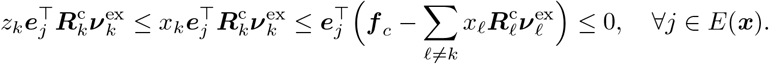

Since the constraint 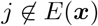 are slack, it follows that there exists *z_k_* > *x_k_* such that

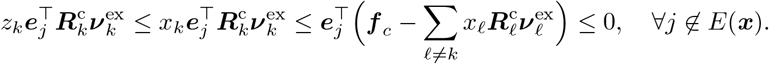

Thus, it follows that (***π**_k_*, 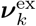) is feasible for (8) when the biomass of the *k*-th species is set *z_k_* > *x_k_*. Therefore, the optimal value when the biomasses *x_ℓ_* and fluxes 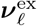 for all species *ℓ* ≠ *k* are fixed, and the biomass of species *k* is set to *z_k_* > *x_k_*, is at least *δ_k_*. Hence, (***x***, ***π***, ***ν***^ex^) is not resource-constrained.

Lemmas 2.5 and 2.6 established the relationship between resource-constrained steady states, GNE for (12), and stability of steady states.

#### Lemma 2.5.

(***x***, ***π***, ***π***^ex^) *is a resource-constrained steady-state GNE if and only if* 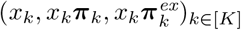 *is a GNE for (12)*.

*Proof.* Suppose (***x***, ***π***, ***ν***^ex^) is a resource-constrained steady-state GNE, but 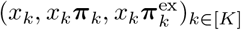 is not an equilibrium for (12). Then there exists a species *k* that can unilaterally improve its solution to (12) i.e. there exists biomass *z_k_* > *x_k_* and fluxes 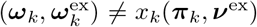 that are feasible for (12). Thus, it follows that 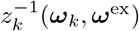 is feasible (8) with an objective value at least *δ_k_*. This contradicts that (***x***, ***π***, ***ν***^ex^) is resource-constrained.

Now suppose (***x***, ***π***, ***π***_ex_) is not a resource-constrained steady state Nash equilibrium. If (***x***, ***π***, ***π***_ex_) is not a steady state GNE, it follows from Lemma 2.3 that 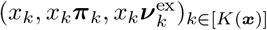 is not a Nash equilibrium for (12). Suppose instead (***x***, ***π***, ***ν***^ex^) is a steady state GNE but it is not resource-constrained. Then there exists a species *k* such that the optimal value of the best response problem (8) with its biomass *z_k_* > *x_k_* is at least *δ_k_*, i.e. there exist feasible fluxes 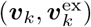 for (8) such that 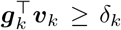. Thus, it follows that 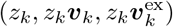 is feasible for (12) with an objective *z_k_* > *z_k_*. A contradiction.

#### Lemma 2.6.

*Suppose* (***x***, ***π***, ***π***_ex_) *is a steady state GNE, but it is not resource-constrained. Then it is unstable to perturbation in the biomass **x***.

*Proof.* Suppose (***x***, ***π***, ***π***_ex_) is not resource-constrained. Then there exists a species *k* and *z_k_* > *x_k_* such that the problem (8) with the biomasses *x_ℓ_* fluxes 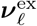 fixed and the biomass of species *k* set to *z_k_* has an optimal solution 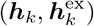 with a value at least *δ_k_*.

Thus, we have that 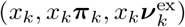 and 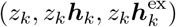 are both feasible for (12). Since the feasible set for (12) is convex, it follows that 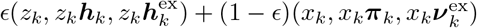 is also feasible for (12) for all *ϵ* ∈ [0,1]. Thus, it follows that the fluxes

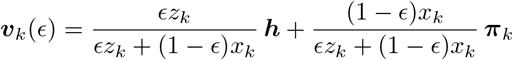

are feasible for (8) with species *k* biomass set to *ϵz_k_* + (1 – *ϵ*)*x_k_* > *x_k_* with an objective value at least *δ_k_*. Therefore, the optimal value is at least *δ_k_*

Consider perturbing the biomasses to 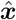 with 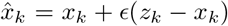 and *x_j_* = *x_j_* for *j* ≠ *k*. Then the result above shows the perturbation persists. Thus, it follows that the steady state GNE (***x***, ***π***, **ν**^ex^) is unstable.

Now we describe the exhaustive search procedure that implies Lemma 2.7. The first step in the search is to write the FBA constraints in terms of the Elementary Flux Modes [37]. Assume that the elementary modes for species *k* are given by

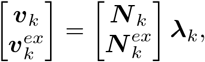

where the mode vector **λ**_*k*_ satisfies **1**^⊤^**λ**_k_ = 1 and **λ**_*k*_ ≥ **0**. Let {(*x_k_*, **λ**_*k*_): *k* ∈[*K*]} denote a Nash equilibrium defined in terms of the Elementary Flux modes. Then the analog of (12) in terms of **λ**_*k*_ is given by

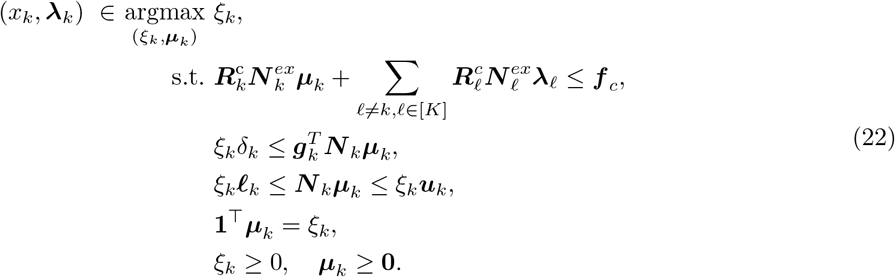

Define the per-species feasible set

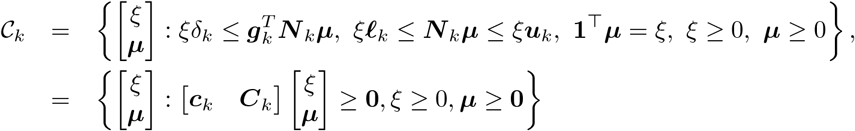

Note that the feasible set 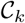 is a cone, and the dual cone 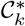 is given by

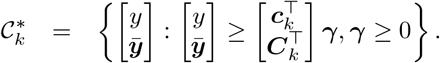

Let 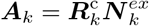 for *k* ∈[*K*]. Then (22) can be reformulated as follows:

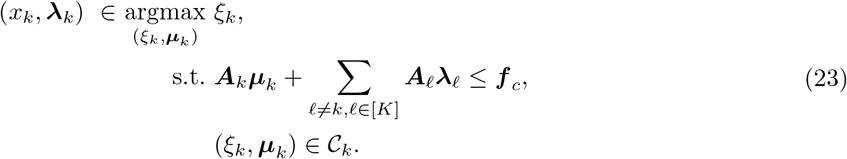

Using duality one can show that (***x***, **λ**) is a steady state GNE if, and only if, there exists *z* = (***z**_k_*)*k*∈[*K*] and ***w*** such that (***x***, ***λ***, ***z***, ***w***) is a solution to

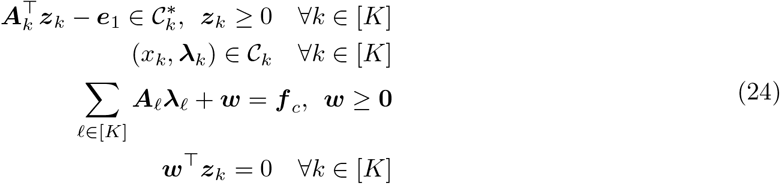

Let *m* denote the dimension of the shared constraints. Then we can enumerate steady state GNE by computing the vertices of the polytopes given by (24) with ***w***[*J*] = **0** and ***z**_k_*[*J^c^*] = **0** for all *k* ∈ [*K*], and doing so for each *J* ⊆ {1,…, *m*}. The following result shows that it is sufficient to only check the stability of such steady states.

#### Lemma A.3.

*Fix J* ⊂ {1,…, *m*}. *Let* (***x***, ***λ***, ***z***, ***w***) *denote a feasible point of* (24) *corresponding to J that is not a vertex. Then it is unstable.*

*Proof.* Let 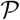 denote the polytope corresponding to *J*. Since (***x***, ***λ***, ***z***, ***w***) is not a vertex of 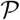,

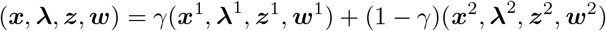

where 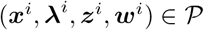, *i* = 1, 2, and *γ* ∈ (0, 1). Thus, for every species *i* it is the case that either 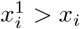 or 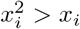.

Fix a species *k*. Without loss of generality, assume 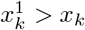. Define

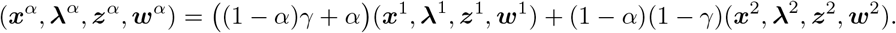

Note that 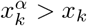 for all *α* > 0, and *x^α^* = *x* + α(*x*^1^ – *x*), and therefore, for all *α* sufficiently small, *x^α^* is a small perturbation of ***x***. Since 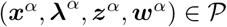, it follows that (***x**^α^*, ***λ**^α^*) is a GNE for (12), and therefore, by Lemma 2.3, (***x**^α^*, ***π**^α^*) is a steady state GNE for (8), where ***π**^α^* is constructed using ***λ**^α^*. Recall that 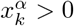 therefore, for ***x**^α^* to be a steady state GNE, we must have that the growth 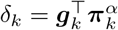. Thus, there exists a GNE such that the state ***x**^α^* does *not* revert back to *x* following a perturbation *p* = *α*(*x*^1^ – *x*).

The vertices of the polytopes given by (24) corresponding to *J* can be enumerated by generating random cost vectors *c* and solving LPs maximizing (***x***, ***λ***, ***z***, ***w***) · *c* subject to (24) and the active sets *J*. Also, in practice one could just search over sets *J* corresponding to metabolites that are externally supplied and/or metabolites that are involved in cross-feeding reactions, rather than all possible subsets of {1,…,*m*} which will involve many community metabolites that are neither supplied nor involved in cross-feeding.

### A.5 Stability to invasion

#### Lemma A.4.

*Suppose a steady-state GNE* (***x***, ***π***, ***π***^ex^) *satisfies the conditions in Lemma A.1. Then, there exists matrices* {***B**_ℓ_*: *ℓ* ∈ *K*[***x***]} *such that*

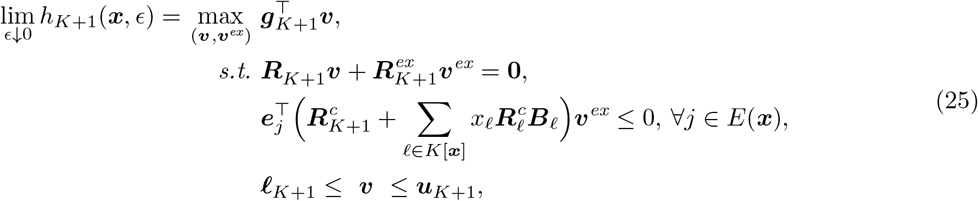

*where* 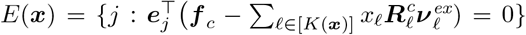, *e_j_ denotes the j-th standard basis vector, and the matrices* {***B**_ℓ_*: *ℓ* ∈ *K*[***x***]} *are defined in the proof below.*

*Proof.* Consider the LP (8). Convert this LP to standard form by adding slacks ***s**_k_* ≥ 0 to the exchange constraints, and splitting the free variables 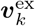 into the difference 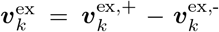, of non-negative variables 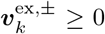. Let *F_k_*(***x***) = {*i*: *i* ∉ *U_k_*(***x***) ∪ *L_k_*(***x***)} denote components of the vector ***π*** that are *not* fixed to their upper or lower bounds, 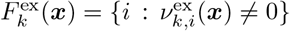 the set of non-zero exchange reactions, and *E*(***x***) the set of active exchange reaction constraints in (8). Then, for all small enough perturbations Δ***f**_c_*, the change in the optimal solution 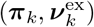 is given by the solution to the following linear equations:

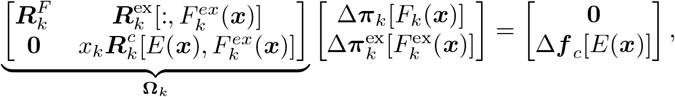

where 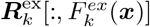 denotes the submatrix obtained by keeping all the rows and only the columns in 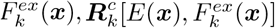 denote the submatrix obtained by taking the rows in *E*(***x***) and the columns in 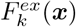, and 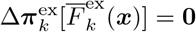, where 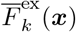 denotes the complement of the set 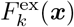. Consequently,

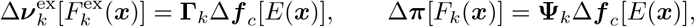

where the matrices **Γ**_*k*_ and **Ψ**_*k*_ are appropriately defined submatrices of **Ω**^−1^. This relationship is valid only if the following constraints hold

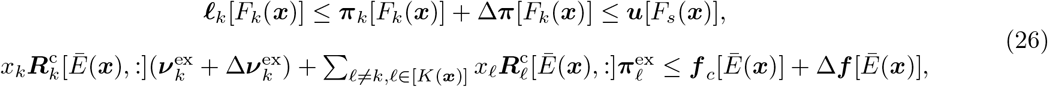

where 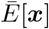 denotes the complement of the set *E*(***x***). Note that these constraints hold as long as the norm of the disturbance Δ*f_c_* is sufficiently small.

The effective change in the RHS for species *k* is given by

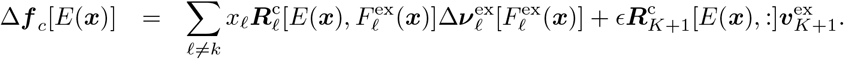

Thus, for all *k* ∈ *K*[***x***], we have that

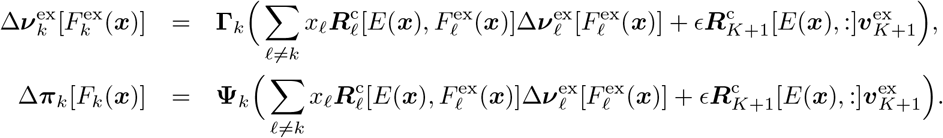

Thus, the perturbation can be written as 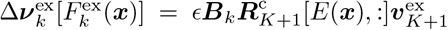, and 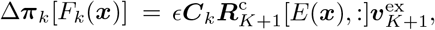, where (***B**_k_*, ***C**_k_*), *k* ∈ *K*[***x***], are a function of (**Γ**_*k*_, **Ψ**_*k*_), *k* ∈ *K*[***x***]. Thus, for all small enough *ϵ*, 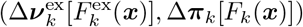 will satisfy (26) for all *k* ∈ *K*[***x***]. More formally, let 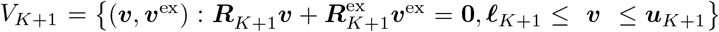. By Assumption 1, we have that the set *V* _*K*+1_ is bounded. Define

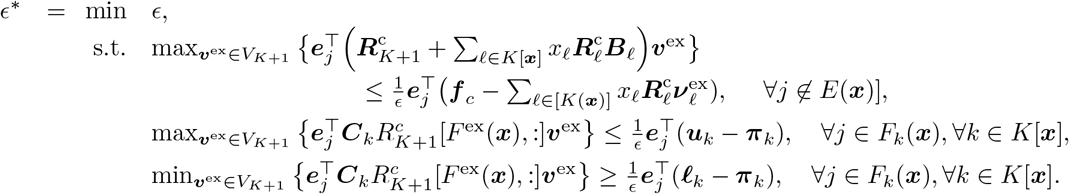

From the definition of the sets *E*(***x***) and *F_k_*(***x***), and Assumption 1, it follows that *ϵ*\* > 0. The contraints corresponding to *j* ∉ *E*(*x*) are slack for all *v*^ex^ ∈ *V*_*K*+1_ for *ϵ* < *ϵ*\*. Consequently, it follows that

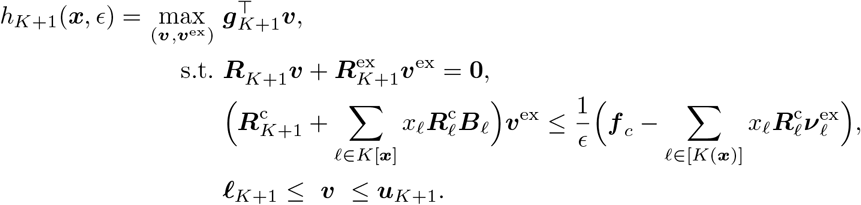

for all *ϵ* < *ϵ*\*. The result follows by taking the limit *ϵ* ↘ 0.

We now address the assumption in Lemma A.1 and Lemma A.4 that the optimal dual vector corresponding to the LP (8) is unique. Note that LPs with continuous parameters [27] have unique dual solutions with very high probability; therefore, it is unlikely that we will encounter problems where the optimal dual solution is non-unique. We can further increase the probability of an optimal dual solution by removing redundant constraints and variables. And, if degeneracy is still encountered, the metabolite supplies ***f**_c_* and reaction bounds ***ℓ**_k_*, ***u**_k_* can be slightly perturbed to generate a new problem with a non-degenerate optimal solution, and such a perturbation is justified since reaction rates and metabolite supply are inherently noisy. Assuming uniqueness of the dual solutions is also standard in work related to differentiating through optimization problems [1, 2], which is exactly what we have done here and in Section A.3.

## Footnotes

1 Note that the same metabolite may be indexed differently in two different species and the compartment.

